# The Role of Attention in Immersion: The Two–Competitor Model

**DOI:** 10.1101/2023.07.10.548435

**Authors:** Daniel J. Strauss, Alexander L. Francis, Jonas Vibell, Farah I. Corona–Strauss

## Abstract

Currently, we face an exponentially increasing interest in immersion, especially sensory–driven immersion, mainly due to the rapid development of ideas and business models centered around a digital virtual universe as well as the increasing availability of affordable immersive technologies for education, communication, and entertainment. However, a clear definition of ‘immersion’, in terms of established neurocognitive concepts and measurable properties, remains elusive, slowing research on the human side of immersive interfaces.

To address this problem, we propose a conceptual, taxonomic model of attention in immersion. We argue (a) modeling immersion theoretically as well as studying immersion experimentally requires a detailed characterization of the role of attention in immersion, even though (b) attention, while necessary, cannot be a sufficient condition for defining immersion. Our broader goal is to characterize immersion in terms that will be compatible with established psychophysiolgical measures that could then in principle be used for the assessment and eventually the optimization of an immersive experience. We start from the perspective that immersion requires the projection of attention to an induced reality, and build on accepted taxonomies of different modes of attention for the development of our two–competitor model. The two–competitor model allows for a quantitative implementation and has an easy graphical interpretation. It helps to highlight the important link between different modes of attention and affect in studying immersion.

## 1 Introduction

Sensory–driven immersion is becoming more and more a topic of our everyday life. The reasons for this are manifold: The rapid development and availability of ‘immersive technologies’; from virtual and mixed reality headsets to virtual audio and haptic displays, and the concomitant development of business models organized around a digital virtual universe (metaverse) [1, 2, 3] but also, more recently, the greatly increased use of digital environments for communication and education in the COVID–19 pandemic see, e.g., [4]. The nature of immersion is addressed in several scientific domains, ranging from computer game research [5], education [6] and entertainment [7] to tracts in philosophy [8]. Even though some categorizations have been proposed such as perceptual [9], sensory [5], and narrative immersion [10, 11] and their subcategories of emotional, spatial, and temporal immersion (see Ryan [10]), science has not really kept pace with the inflationary use of the catch–all term ‘immersion’ by defining what ersion as generic concept actually is; a problem which has been pointed out by several researchers, e.g., see McMahan [12], Chasid [13], Nilsson et al. [14], Murray [15], Agrawal et al. [16]. Defining immersion is not just important from a theoretical perspective. It also has practical implications. Immersion is a state that does not seem to stand up well to overt introspection in the moment. It is not useful to ask someone “are you immersed?” any more than it is useful to ask “are you asleep?” Immersion seems to rely at least in part on a sort of self–delusion, such that the mere conscious awareness of the sense of being immersed may be sufficient to destroy it. Thus, if we are to develop methods to evaluate whether or to what degree a person has become successfully immersed in an induced reality, whether one induced by reading a book or by the latest headset based virtual environment, those methods must necessarily be covert and unobtrusive, and applicable without interfering with the immersive state itself.

Methods of evaluating immersion must focus on attention. In a recent, comprehensive attempt to define immersion in the context of audiovisual experiences, Agrawal et al. [16] defined immersion as “the state of deep mental involvement in which the subject may experience disassociation from the awareness of the physical world due to a shift in their attentional state”. In addition to Agrawal et al. [16], several other scientists have highlighted the prominent role of attention in immersion, e.g., see [15, 11, 17, 18], linking research on immersion to established concepts in cognitive neuroscience. Here, we start from an attentional emphasis similar to that of Agrawal et al. [16], but build out details of the model with the goal of using predictions from existing cognitive neuroscience models of attention to guide research on immersion. For example, the assumption that immersion requires the projection of attention to an induced reality would suggest that processing resources must be allocated (see Kahneman [19]) to information associated with that reality, and that physiological measures associated with attentional allocation could best be employed to assess immersion as well.

## 2 Different Modes of Attention in Immersion

In order to model the attentional properties important for understand immersion, we similarly begin with the idea that immersion requires being focused on a task in such a way that one loses awareness of events or sensations outside of it [16]. The increased awareness of one stimulus (or set of stimuli) at the expense of others is, of course, a hallmark of simple selective attention. Thus this preliminary characterization of immersion could, in principle, be accommodated entirely in terms of models developed to account for selective attention and ‘inattentional blindness’ [20, 21, 22]. Nevertheless, there are surely other qualities besides selective attention that must be present when one is truly immersed in an induced reality. Evidence that many researchers are aware of this additional quality can be found in literature addressing the importance of distinguishing immersion from related terms used mainly in virtual reality research, in particular, presence [23, 24, 25, 26], transportation [27, 28], flow [26, 29], and envelopment [30], see Agrawal et al. [16] for detailed discussions. It is beyond the scope of this article to evaluate the degree to which immersion does or does not differ from these various other qualities; the authors in [16] did already an excellent analysis of these differentiations. However, it is important to note that one factor that is consistently cited as distinguishing immersion from simple selective attention is the importance of self-initiation and intrinsic motivation in immersion, see Deci and Ryan [31], Ryan and Deci [32], Richter [33], Kurzban [34], Herrmann and Johnsrude [35].

We suggest that the concept of intrinsic motivation is particularly relevant here, such that immersion is strengthened when the motivation to attend to an induced reality is supported specifically by task- or narrative-related properties of the activities being conducted in that reality. That is, there seems to be more to immersion than simply the intentional, self-initiated direction of attention toward the sensory streams of that reality. Immersion requires more than just the will or the intention to attend to some stimuli more than others, it requires the intention to attend to some stimuli more than others. We propose that immersion requires attention to be directed to mental representations (memories, narrative structures, schemas, etc.) associated with a particular set of coordinated stimuli existing within the induced reality because those stimuli *fit together* in some particularly compelling way.

Indeed, if we are to develop a model of attention in immersion that can be applied broadly to a wide variety of immersive situations, we must also consider circumstances in which one becomes immersed in a wholly internal reality, as in “daydreaming.” Even in the absence of any novel sensory input, it is still possible to become immersed in an induced reality that consists entirely of purely self–generated internal objects and that therefore depends to an extreme degree on the deployment of so-called ‘internal attention’ [36, 37]. Such cases highlight the first of two important distinctions between kinds of attention. Internal attention refers to attention directed toward internally generated mental representations such as schemas and narratives. In such extreme cases, internally directed attention is all there is to support immersion. In the technologically more dominant case of sensory–driven immersion – whether as simple as visual input from reading a electronic book or as advanced as using multisensory virtual, augmented, or mixed reality high–fidelity technologies – the induced reality may be established by immersion–provoking sensory events but we argue that the sense of immersion in that reality still likely requires the development, maintenance, and attention to internal representations as well.

### Attention in Sensory Dynamics

Therefore, to cover the sensory dynamics when dealing with a sensory–driven reality as well as the internal processes related to the narrative and schemas of the induced world e.g., the unfolding of the story [10] (and potentially also the self-generated/imagined sensory inputs of an internally-generated world, e.g., from daydreaming or reading), our model employs two different but established taxonomies of attention. These attention taxonomies are integrated in a conceptual model that also incorporates affect, effort and motivation to pursue goals in an induced reality (in line with Deci and Ryan [31], Ryan and Deci [32]) enabling future consideration of the role of narrative engagement (in line with Albrecht and O’Brien [38], Green and Brock [39], Bilandzic and Brusselle [40]).

Although we have already mentioned the importance of internally directed attention in maintaining a sense of immersion, when discussing sensory dynamics it is simpler to start with the traditional taxonomy of exogenous and endogenous of attention in sensory processing, see Müller and Rabbitt [41], Spence and Driver [42], Berger et al. [43], Sani et al. [44], Jigo et al. [45], Ren et al. [46]. Exogenous or bottom–up attention to the respective sensory modality is driven by the physical properties and saliency of the stimuli. Such properties might include location, intensity, and abruptness of onset in any domain, as well as more sense-specific properties such as tonality or roughness in the auditory domain and color or brightness in the visual domain. It is important to note that these properties should affect the distribution of attention irrespective of whether the stimulus to which they are attributed is perceived in the actual or induced reality. Exogenous attention is involuntary and comparatively automatic. In contrast, endogenous attention refers to a (goal–driven) top–down attention which is voluntarily allocated to an object or stream of interest. Endogenous attention is closely linked to intention, motivation and thus goal pursuit in a broader sense, e.g., see Deci and Ryan [31]. Which sensory information stream is within the attentional focus thus depends on a combination of exogenous and endogenous factors such as might be modeled by a probabilistic stream selection model [47]. Here different exogenous and endogenous weights are assigned to the individual sensory streams and define their probability for being selected, i.e., being within the attentional focus. This probabilistic selection scheme can be seen as computational approaches to the biased competition model which is well-known in visual perception in Desimone and Duncan [48] and has also been discussed for the auditory modality by Shinn-Cunningham [49].

The exogenous and endogenous components of the model reduce the selection of sensory streams in an actual or induced reality to the abstract notation of exogenous and endogenous weights. However, to understand the role of attention in immersion, and, in particular, to understand what drives attention to induced realities, we need to have a closer look at the concepts that define these weights. For this, we apply the taxonomy of Chun et al. [36] of internal and external attention which is based on the types of information that attention operates over. External attention is associated with the selection and modulation of sensory information, i.e., information originating from outside the observer. It can thus be thought of as similar in some ways to exogenous attention, but, importantly, can also be endogenously directed. For example, external attention may be directed by endogenous systems when directing attention to a particular location in an otherwise static visual scene, e.g., when waiting for a stoplight to change color. In this case, the observer is directing the distribution of attention – this is endogenous attention – but it is being directed toward an external stimulus. Internal attention, in contrast, refers to the selection, modulation, and maintenance of internally generated information such as task rules, schemas, responses, and elements of long–term or working memory [36]. It is this internal attention to narrative structures, schemas, and mental representations related to the coherence of the induced reality that we have already identified as being so crucial for maintaining the state of immersion.

Even though the exogenous/endogenous and internal/external modes of attention are related, see Chun et al. [36], they must be distinguished if we are to map out the role of different processes when paying attention to an induced reality, especially in a conceptual, let alone a computational implementation. Whereas the exogenous/endogenous taxonomy directly reflects the sensory dynamics of the competition between the information present in each of the two realities, the internal/external taxonomy is more relevant to understanding the importance of narrative and conceptual factors such as the unfolding of a story line while reading a book, the ‘willing suspension of disbelief’ when interacting in a world with, perhaps, unrealistic physics or events, and even the case of daydreaming in the absence of any sensory input. Thus internal attention is closely linked to ‘being in harmony with the induced reality’ and goal pursuit, e.g., see [31] and provides, even in sensory–driven immersion, a framework to analyze higher– order, i.e., not just reflexive, interactions with affect as we will see below.

The “what” and “why” of goal pursuits in humans is a long lasting debate, see [31]. Why somebody should seek goals in an induced reality is certainly highly individual and context sensitive. The prominent role of the narrative in immersion has been stressed by several researchers, e.g., see [10, 50, 5, 11]. More accessible than a goal itself in our analysis, is the motivation in pursuing this goal and the acceptance of the narrative. We assume that paying attention to the induced reality has to be rewarding (in a broader sense, i.e., in terms of achieving a goal, whether that is to have fun, gain information, communicate with others, etc.). However, we assume that there is always a competition between the induced and the actual reality as the latter has an inherent personal relevance. From a purely endogenous point of view, if there is no ongoing motivation to pursue a goal in the induced reality, the attentional focus must sooner or later shift back to the actual reality as sensations from that reality begin to intrude (if nothing else, interoceptive sensations related to posture, hunger, thirst, fatigue, etc.). Exogenous processing must also play a role, particularly in terms of reflexive attention shifts. For instance, aversive and abrupt sound stimuli that immediately capture our attention seem likely override voluntary attentional processes and their respective endogenous weights, respectively.

We have previously considered the importance of exogenous attentional capture in the context of distraction and annoyance from distracting sounds [51]. Indeed, distraction seems like a reasonable characterization of the involuntary shift of attention from the induced reality stream(s) back to actual reality under the influence of a powerful real-world stimulus such as the sound of a door slamming or perhaps the interoceptive feeling of hunger or thirst. It is thus also possible to model the reciprocal case in the same way, such that when a stimulus from the induced reality is sufficiently strong to draw attention (back) toward that domain, it acts as a ‘distractor’ from the real world. However, it seems likely that such purely exogenous shifts of attention might serve mainly to interrupt immersion and would be unlikely to result in immersion on their own. In order to support immersion in an induced reality, a set of stimuli must be sufficiently compelling to hold the focus of attention to the exclusion of stimuli from outside that reality, and we would argue that for the most part this would require at least some degree of endogenous attention to (at least) some internal representations associated with that reality. Indeed, we can conceive of cases in which one’s attention is drawn back again and again to an immersive experience, for example when one cannot stop thinking about a compelling novel or engrossing film. Even though one might argue that such cases represent something other that true immersion, such as perhaps presence or transportation, if we do think of them as representative of immersion to some degree then such cases, which seem typically to involve a high degree of attention to internal representations, especially narrative structures and conceptual schemas, further support our argument that immersion depends heavily on attention to internal representations. In summary: in our two–competitor model, sensory streams emanating from the induced reality compete with those from the actual reality, i.e., the physical world around us and inside us, for the allocation of our attention. The induced and the actual reality are represented as orthogonal axes in a two dimensional plane in which an attentional focus vector is mapped. As the model employs established taxonomies in neuroscience, it is capable of serving both to inform the design and conduct of future research on immersion, while also enabling a direct translation of objective psychophysiological measurement techniques to immersion research.

## 3 Conceptual Taxonomic Model

In this section, we establish a conceptual taxonomic model to study immersion using the aforementioned concepts. To do so, we introduce a simple mathematical framework which might support future experimental designs by quantitative means. We start with an attentional focus vector **a** *∈* R^2^ (we use bold letters for vectors) in a two dimensional plane with the actual reality and the induced reality as different dimensions, see Fig 1 (a). Orthogonality assures that the mapping to the induced vs. the actual reality are mutually exclusive. It is worth to emphasize that this ‘attentional focus vector’ is a highly simplified mathematical construct, see Sec. 4 for details. It is simply used to provide a continuous (smooth) transition between the two–competing dimensions of resource allocation. Immersion requires now the projection of the attentional focus vector on the induced reality axis, i.e., the component of **a** which is represented on the ir–axis. We denote this component by **a**_ir_. Along this line, we denote the projection of the attentional focus vector to the actual reality, i.e., the ar–axis, by **a**_ar_, see Fig. 1 (a). So we have that **a** = **a**_ar_ + **a**_ir_. Note that this can be written as **a** = *a*_ar_**e**_1_ + *a*_ir_**e**_2_, where *a*_ar_ and *a*_ir_ is the magnitude (or length) of **a**_ar_ and **a**_ir_, respectively. The vectors **e**_1_ and **e**_2_ are the standard basis vectors of a two dimensional Cartesian coordinate system defining the competing directions of the actual and the induced reality. In a bistable perception, i.e., either you perceive the induced or the actual reality (see blue/green transition at *φ* = *π/*4 in Fig. 1 (left)), the model can also map a distraction framework (e.g., see McRae et al. [52]). For instance, for the assessment of ‘immersion’ in the induced reality, the projection *a*_ar_ represents the amount of distraction from the induced reality.

**Figure 1:**
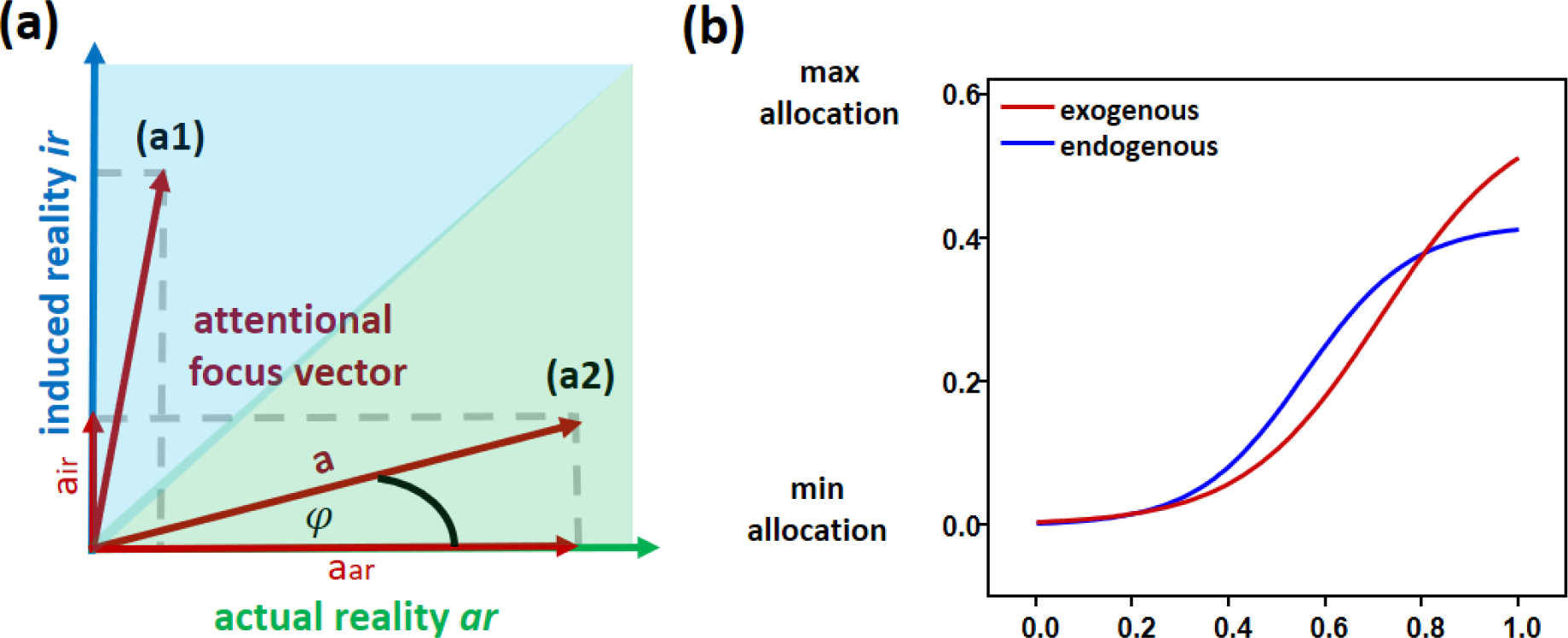
(a) The two–competitor model along the orthogonal axes of the actual and the induced reality. In state (a1) the projection of the attentional focus vector is mainly on the induced reality which ‘wins’ the competition; a necessary condition for a high ‘immersion’. In state (a2) the attention is mainly on the actual reality making it the ‘winner’ of the competition. The decomposition of the attentional focus vector in its component *a_ar_* and *a_ir_* is also shown. (b) The transfer function Ω*_α,β,γ_*(*x*) for *x ∈* [0, 1.2] and *α* = 0.58*, β* = 7*, γ* = 5 (exogenous) and *α* = 0.42*, β* = 9*, γ* = 5 (endogenous).

The model represents how the attentional focus varies between the competing dimensions. The dominant stream in the individual reality is the momentary “winner” of a stream selection based on the exogenous and endogenous weights to several input streams. A probabilistic stream selection for this is described in [47] but an alternative is suggested below. In module 1 below, we will first have a closer look at the dynamics of the attentional focus between these “winners” or dominant streams in the individual reality and their associated weights.

### 3.1 Module 1: Exogenous and Endogenous Sensory Dynamics

The direction of the attentional focus vector **a** is now given by the exogenous and endogenous weights of these dominant streams in the individual realities. In dynamic regimes, the vectors as well as the weights depend on time *t ∈* R. We denote the exogenous and endogenous weights in R by 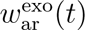 and 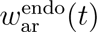 for the actual reality and by 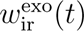 and 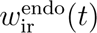 for the induced reality. We define a transfer function 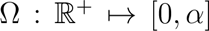 to map the (possibly not normalized) weights (see below) to [0*, α*], where *α ∈* R*_>_*_0_ is the maximum resource allocation. Here we use the sigmoid function 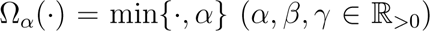 but other possible choice are, for instance, Ω*_α,β_*(*·*) = *α* tanh(*β·*) or Ω*_α_*(*·*) = min*{·, α}* (*α, β, γ ∈* R*_>_*_0_). Here *β* and *γ* determine the transfer sensitivity of the mapping, see Fig. 1 (b).

There is a long-standing debate in the literature over how the exogenous and endogenous mechanisms of attentional allocation interact, especially across modalities, e.g., see Müller and Rabbitt [41], Spence and Driver [42], Berger et al. [43], Sani et al. [44], Jigo et al. [45], Ren et al. [46]. Some models suggest that there should be a stronger interaction (i.e. greater weights in our model) under high demand due to a competition for resources, see Berger et al. [43]. While we make explicit choices here for the purpose of developing a working computational model, we acknowledge that further research is likely to suggest the need for further refinements in this regard. For example, due the evolutionary significance for survival of the auditory modality for automatic orienting in primates (e.g., Carretíe [53], Strauss et al. [54], see also “Advantages & Costs of Front–Facing Eyes” in Allman [55]), we give priority to the exogenous channel to ‘override’ endogenous weights in extreme conditions, e.g., due to abrupt, very intense sounds. However, this weighting could be modified depending on the specific method for inducing immersion, i.e. sound-damping headphones, goggles, etc. For the present model, we use different transfer functions Ω*_α,β,γ_*(*·*) for the exogenous and endogenous channels, see Fig. 1 (b) for examples. When using the notation *α^′^, β^′^, γ^′^* for the parameters of the endogenous transfer function, the individual attentional projections are given by

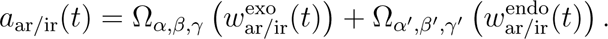

Using these projections, the magnitude (or length) of the attentional focus **a**(*t*) is given by 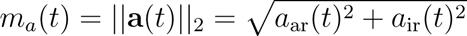 with the corresponding angle

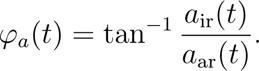

Then, in this two–competitor model, *φ_a_* defines the projection of attention to the induced reality, i.e., fully focused for *φ_a_* = *π/*2 and not focused at all for *φ_a_* = 0.

### 3.2 Module 2: External vs. Internal Attention and Affect

In order to assign the weights 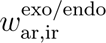 to sensory (or internally generated) streams or objects, we employ a conceptual link inspired by the taxonomy of internal versus external attention of Chun et al. [36] as a more (psychologically) representative alternative to the probabilistic stream selection in [47]. The demands for a generally valid model of these attention modes in immersion are of course immense as, especially regarding internal attention, they crucially depend on many factors including the induced reality type (computer games, educational settings, virtual meetings, entertainment etc.), the nature and coherence of the narrative, the participant’s personal interest, and several other individual factors. As a first step, we suggest an orthogonalization of these attention modes which we have used in Strauss and Francis [37] to analyze effortful listening. Here the resulting attentional vector is used to define an (attentional) effort response in the sense of Sarter et al. [56]. This conceptual model is then further coupled with affect in a closed–loop system that includes related modules such as motivation and reward by Schneider et al. [57] to represent cognitive fatigue effects in coordination with the effort response. The interaction of resource allocation and affect was also carefully analyzed in Francis and Love [51], Herrmann and Johnsrude [35]. Models of this type are not necessarily restricted to effortful tasks in the induced reality and can be generalized to provide an endogenous response reflecting affect related factors. In fact, when using a proper normalization, this affect-dependent endogenous response may directly define, at least in a first approximation, the weights 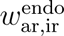 in Module 1 as described earlier.

For instance, when losing the motivation to solve an effortful task in the induced reality (e.g., the difficulty of an ‘immersive’ computer game is too high, resulting a bad effort/reward ratio), the endogenous weight will decrease over time. But the very same decrease in motivation happens when watching a boring movie over a longer time, even though this task not necessarily effortful. For the sake of simplicity, for the moment we represent models of this type by a multivariate map 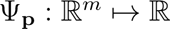, representing the endogenous response to *m* sensory or internally generated streams or objects, denoted by **s**(*t*) = (*s*_1_(*t*)*, s*_2_(*t*)*, …, s_m_*(*t*))*^T^*, and specified by a set of parameters **p** *∈* R*^n^*. Thus, the weights 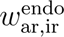 are simply given by

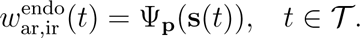

Here *T* denotes the interval of the simulation time. Also a multisensory integration (see Sec. 4) may be included in this function (or module) as a missing multisensory fusion or a lack of concurrency of the sensory inputs (see Calvert et al. [58]). Such a feature might reflect the damage to immersion that might be caused by a low–fidelity system that generates a (multisensory) induced reality in which the different sensory streams are poorly coordinated in time (think of a poorly dubbed movie in which the movement of the lips does not line up well with the sounds of the dubbed speech). Such mis-synchrony may well reduce the feeling of presence [14] or other quality important for the immersive experience, see the discussions in Marucci et al. [59], and would be reflected in a loss of endogenous attention directed toward internal mental representations (schemas, narrative structures, etc.) associated with the coherence of the reality, that is as smaller endogenous weight 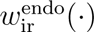, see Sec. 4.3 for further discussions.

To model the exogenous weight, we define the parameteized, multivariate function 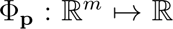. It reflects the attention directed toward sensory stimuli based on their perceptual and affective salience, e.g., brightness, loudness, valence and arousal. Higher order features, e.g., computationally extracted voice features [60, 61] or facial features [62] in case of overt attention can also be integrated in the model Φ**_p_** (see Section 4). Thus we have that

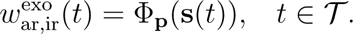

The entire two-competitor model with these maps and the sensory input is shown as schematic diagram in Fig. 2.

**Figure 2:**
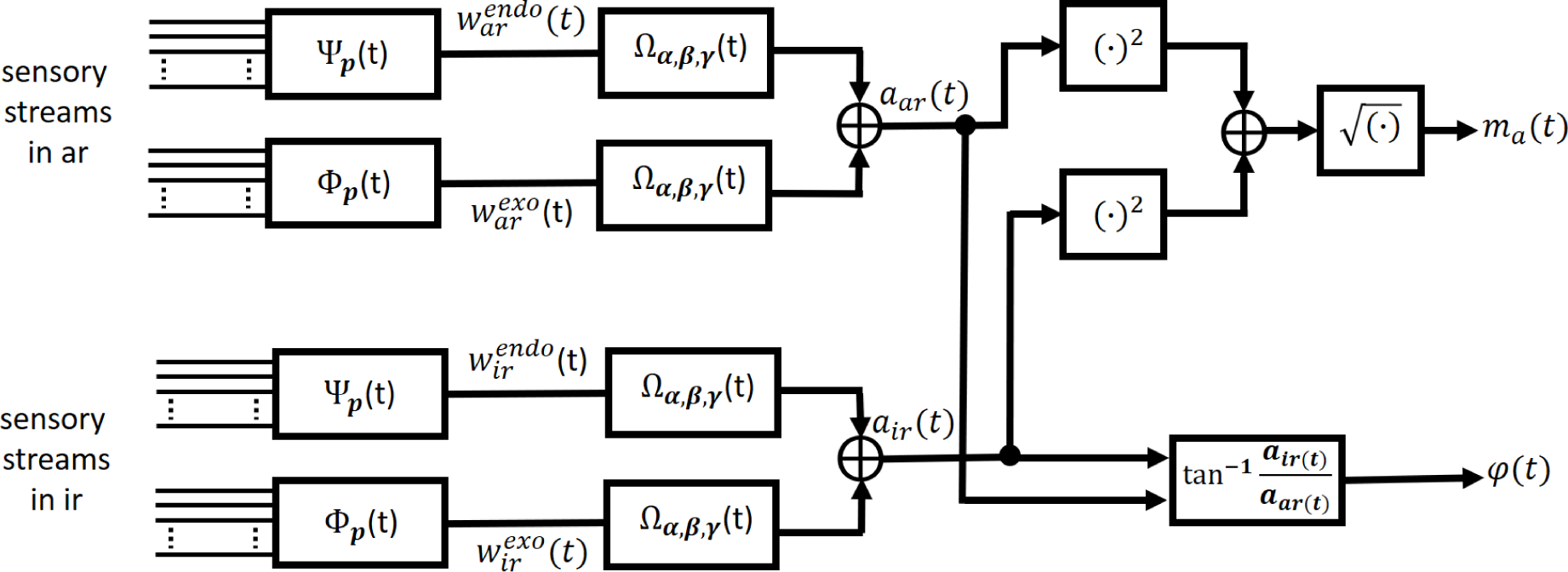
A schematic diagram of the dynamic two–competitor model. The input is given by the sensory streams as well as defined by the parameters **p** of the functions Φ**_p_** and Ψ**_p_**. The output is the angle *ϕ*(*t*) representing how much attention is projected to the induced reality. The magnitude of the attentional focus vector is given *m_a_*(*t*).

#### Exemplary Implementation of an Effortful, Fatiguing Task in the Induced Reality

As an example, we present an implementation of an effortful task in the induced reality such as playing a computer game with an increasing level of difficulty. For this, we focus on the map Ψ**_p_**(*t*) which is implemented using the fatigue model in Schneider et al. [57] in which the sensory input streams **s** are represented directly in terms of their internal and external demands^1^. The effort response to these demands [37, 57] defines the endogenous weight 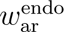 whereas the other weights are modeled by simple oscillatory functions in time, see Fig. 3. In 3 (1. row) different (actual) motivation profiles are shown. On the left, playing the game is rewarding (good reward to effort ratio [57]) and we have a rather constant motivation over the 20min time of playing. On the right–hand side, the skills of the player cannot catch–up with the difficulty, resulting in a bad reward to effort ratio, fatigue, and thus a decease in actual motivation over time up to the point of giving up completely in the end. The resulting endogenous weight mapped by Ω*_α,β,γ_*(*·*) is shown below (blue line) along with the other mapped weights using simple oscillatory functions as model. Note that around 10min there is a short intense exogenous activation in the actual reality reflecting a sudden, very loud sound such as a slamming door (black line). The endogenous weights directed toward the actual reality are moving up over the time in this model because we assume that there is necessarily increasing pressure to do something else in the actual reality, if only due to the biological necessity of survival. The increasing difficulty in the game is, in contrast, associated with a more rapid exogenous stimulation in the induced reality which is modeled by the (up–)chirp function. The resulting magnitude *m_a_*(*t*) and angle *φ_a_*(*t*) are shown below. It is noticeable that *φ_a_* reflects the competition between the individual realities for the allocation of attention.

**Figure 3:**
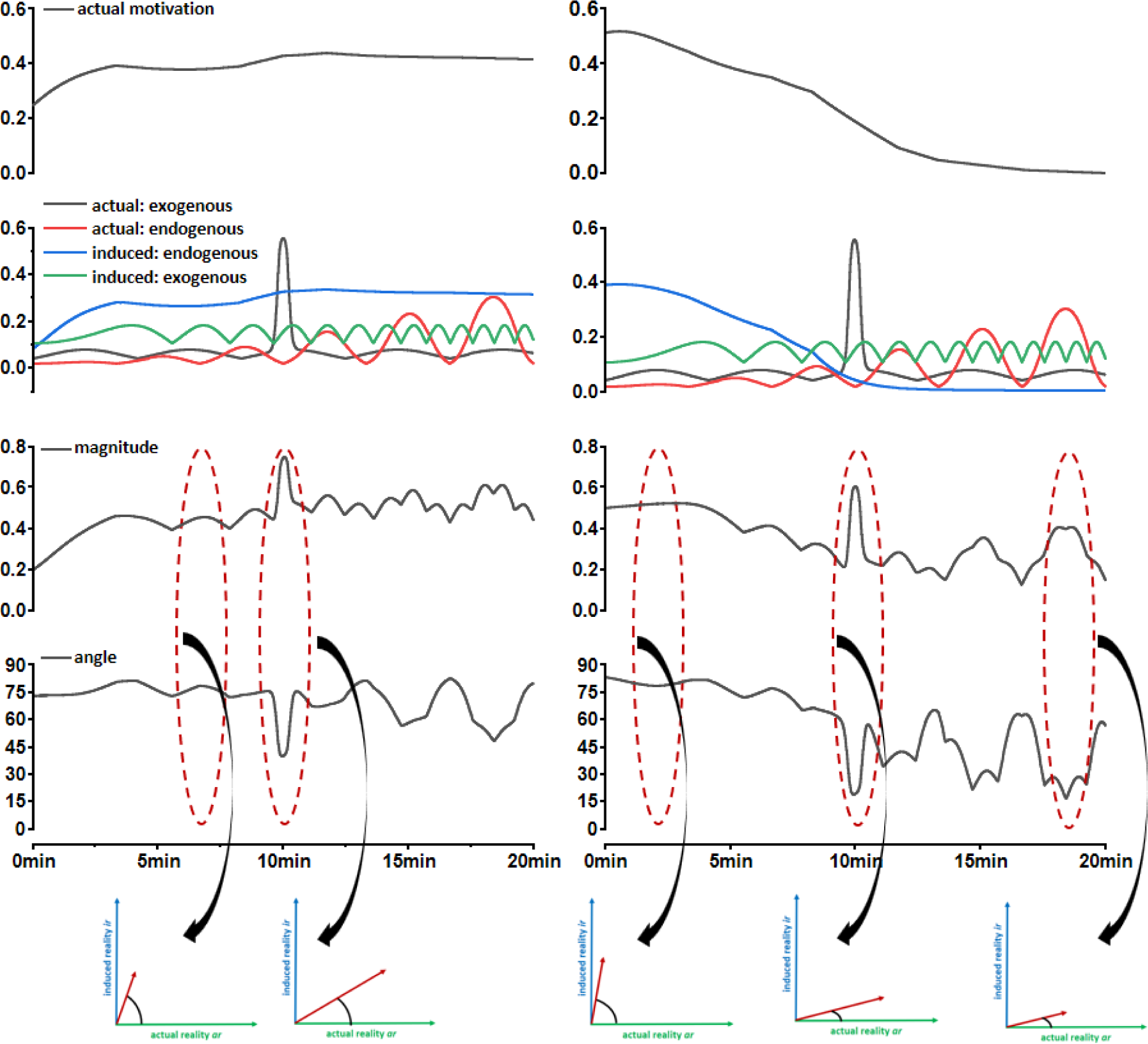
The first row (top) shows the actual motivation profile for a rewarding activity (left) and a very effortful fatiguing activity (right). Second row: The weights mapped by Ω*_α,β,γ_*(*·*); here 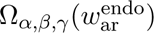 stems from the actual motivation above whereas the others are modeled by simple oscillatory functions. Third row: The magnitude of the attentional focus vector *m_a_*(*t*). Fourth row: The angle *φ_a_*(*t*) of the attentional focus vector. Beside the angles *φ_a_*(*t*) in degrees, all units are normalized such that 1.0 is the maximum allocation.

#### The quantitative model in summary

Module 1 defines the projection of the attentional focus vector to the actual and induced reality by accumulating the exogeneous and endogenous weights to stimuli in the respective reality. These weights are proportional to the attentional allocation in the respective reality. The resulting two projections, i.e., to the actual and induced reality, define the attentional focus vector and the corresponding angle between this vector and the respective reality. The closer this angle is to 90*^◦^* (which is *π/*2 in radians), the maximum angle in the defined setting, the more attention is allocated to the induced reality. So, everything depends on the weights. The weights are defined by module 2 using the taxonomy of internal and external attention. In fact, we defined a (generic) map which maps the sensory streams in the respective reality to the exogeneous and endogenous weights using the internal and external mode of attention. As presented, this map can be any model which provides this, though an explicit, generally valid formulation is beyond the scope of this paper. However, by employing a model of cognitive fatigue which links internal and external attention to reward and motivation (i.e., affect-related quantities) from [57], we provided an example for such a map.

## 4 Discussion

Although preliminary in terms of detail, the present model provides a conceptual framework linking the neuroscience of attention to fundamental and applied immersion research. As such, it should provoke new, more specific research questions related to the role attention in immersion and its interaction with affect. In addition, we hope that it also stimulates researchers in cognitive and affective neuroscience to apply their knowledge to immersion research. To that end, in this section we present some preliminary observations from the model that suggest some potential avenues for future research.

### 4.1 Optimizing Immersion in Induced Realities: Internal Attention, Motivation, and Affect

In this section, we outline some ways in which the present model might contribute to studying the optimization of immersion; an important problem particularly in the development of more effective immersive technologies.

For example, the presented two–competitor model maps the role of attention in immersion using two distinct but related taxonomies of attention, and these must be considered together when optimizing immersion. To optimize immersion, the angle *φ_a_*(*t*) *∈* [0*, π/*2] has to be maximized in the two–competitor model. Conceptually, the model was designed such that it can in principle cover the full range of examples of immersion discussed in the literature, from being ‘immersed’ in a daydream or in reading a book to the more commonly considered case of multimodal immersion in the metaverse projected using high fidelity audiovisual technology. Reflecting the diversity of these examples in terms of the diversity of attentional systems, this model is capable of achieving angles close to *π/*2 (i.e., 90*^◦^*) (representing a very high degree of immersion) in very different ways. In the purely narrative driven case of reading a book, an interesting narrative absorbs your internal resources and creates an induced reality, e.g., by unfolding a story in your mind. In the technology driven case, a highly detailed array of multisensory information stimulates a complex interplay of internal attention to the narrative and external attention to multiple sensory channels, driving a highly dynamic exogenous and endogenous allocation of attention. Whereas in the former case, internal attention and the associated interplay with motivation and affect drives immersion, in the latter case, exogeneous input and external attention play a crucial role as well. In summary, both the amount of attention (length of the arrow) that is being directed, and also the direction (angle) it is distributed toward, contribute to the degree of immersion, but equivalent values of these measures can in principle be achieved through different mechanisms.

In many ways, it seems that the technology driven approach would have more flexibility, or perhaps greater capacity, to create immersion, but this may only be superficially the case. Most obviously, modern “virtual reality” immersive systems can attract attention exogenously through visual, auditory, and even tactile stimulation simultaneously (though this may have disadvantages as well, see below). More generally, effects such as abrupt onsets of emotionally loaded stimuli may represent an excellent exogenous means to pull and keep someone’s attention to an induced reality – not an unknown concept in the Hollywood film industry (e.g., the infamous “jump scares” of formulaic horror movies). However, it also seems likely that the effectiveness of such emotional stimuli at maintaining immersion, no matter how good they are at initially attracting attention, will still depend in the long run on their consistency with the narrative and with observers’ expectations, i.e., with mental representations that depend on the endogenous commitment of internal attention, and such narrative consistency can also be achieved through less technological means. Nevertheless, these cases highlight the close relationship between attention and intention [63] or motivation, and thus also emotion [64] and associated psychophysiology.

### 4.2 Objective Assessment of Immersion

In this section, we review and discuss a few issues that arise when considering how we might use objective physiological measures to assess immersion in terms of the attentional and affective properties identified in the presented model. As a myriad of methods can be used for this purpose (e.g., electro– and magnetoencephalography, functional magnetic resonance imaging and near–infrared imaging), we focus mostly on widely available and ecologically valid electrophysiological assessment methods, in particular, electroencephalographic methods in the following.

Induced realities, mainly in the form of (visual) virtual reality concepts, are of increasing interest to researchers intending to enhance the ecological validity of their experimental designs in cognitive neuroscience and affective research, e.g., see Beck et al. [65], Wong et al. [66], Reggente et al. [67], Nicol et al. [68], Marucci et al. [59], Hofmann et al. [69], Yu et al. [70]. However, for such methods to be broadly applicable, it is important for researchers to be able to verify that participants are indeed responding to stimuli in a (relatively) uniformly immersed condition. The assessment of immersion has been addressed by a variety of neuroscientific and neurotechnological measurement methods, e.g., to differentiate between concentration and immersion [71], to quantify the degree of presence [72], or to characterize binaural sound immersion [68]. A very recent review by [18] has analyzed different electroencephalographic methods to assess attention, work load (see [73, 74]), and mental fatigue for possible applications in attention detection in virtual environments, focusing on technologically induced realities. In particular, these authors along with others [75] have highlighted the importance of being able to distinguish between the internal and external modes of attention in virtual and augmented reality environments when using neurotechnological measurements, a task further highlighted by the presented model.

#### Magnitude and Angle of Attentional Focus

The suggested two–competitor model, with its emphasis on distinct taxonomies of attention, highlights the importance of attention allocation between the competing realities but also across the internal/external and exogenous/endogenous dichotomies, distinctions which provide greater rigor for the design of experiments assessing the depth of immersion. In particular, the two–competitor model stresses that beside assessing the magnitude of the described attentional focus vector *m_a_*(*t*) to internal or external processes (related to manifold electrophysiological correlates of attention, e.g., see Näätanen [76], Hopfinger et al. [77], Parasuraman and Wilson [78], Hillyard and Picton [79], Mangun [80], Magosso et al. [81]), it is also important to get the direction of attention represented by the angle *φ_a_*(*t*). A straightforward way to implement this is to assess the attentional factors not in total, i.e., the magnitude *m_a_*(*t*), but rather to specific events in the respective reality, i.e., its components *a*_ar_(*t*) and *a*_ir_(*t*). In other words, if we know about the occurrence of events in a particular reality, we can assess the attention paid to them as well as any affective response(s) associated with them. This requires the co–registration of several measurements, including evoked, event–related, and induced electroencephalographic measures which can be associated to ‘events’ (identifiable fluctuations in ongoing measurements) in the respective reality., e.g. see Pfurtscheller and Lopes da Silva [82], Mangun [80], a methodology that can also be referred to as entrainment assessments (see Lakatos et al. [83], Horton et al. [84]). Electroencephalographic measures of external (selective) attention paid to short auditory, visual, or somatosensory events can be applied for such purposes, e.g., see Hillyard et al. [85], Hansen and Hillyard [86], Hillyard et al. [87], David et al. [88], Lehser et al. [89], Hillyard and Picton [79], Mangun [80]. Within the respective reality, more specific types of attention can be assessed by neuroimaging techniques such as feature based, spatial or temporal attention, see Luck and Hillyard [90], Giesbrecht et al. [91], Olivers and Meeter [92], Hillyard and Picton [79], Mangun [80] to further analyze the attentional dynamics within the modality if necessary. Even incongruities between internally generated patterns/predictions and the attended sensory streams can be assessed with such electroencephalographic methods and oddball paradigms, see, e.g., Kutas and Hillyard [93], Polich [94]; thus such methods are closely linked to internal attention and the narrative processing of the induced reality. Research of this sort is particularly well-represented in the auditory modaltiy, where electroencephalographic temporal response function or stimulus reconstruction methods are frequently used to assess to which speech stream external attention is directed to Mesgarani and Chang [95], Power et al. [96], Wong et al. [97], Schäfer et al. [98]. Recent work has also shown that a non–invasive surface electromyogram of the vestigial auricular muscles can be used to decode the direction of exogeneous and endogenous attention to short sounds and competing speech, see Strauss et al. [54].

While directing selective attention to exogenous events in the induced reality is certainly a significant condition for immersion in the typical sensory–driven “virtual” reality, as we have suggested, internal attention must also play a necessary role. Research on assessment of internal attention has tended to focus on *α*–power [99], but other measures may be useful, including pupillometry [100], and future research should also not rule out combining attention assessments with physiological measures of orienting and affect (see Valenza and Scilingo [101], Marucci et al. [59]) or other autonomic nervous system responses, e.g., see Valenza and Scilingo [101], Francis and Oliver [102], that might help link attention to the emotional qualities associated with the internal representations (narrative structures, schemas, etc.) relevant to the induced reality.

#### Detecting and Annotating Events in “Reality”

In order to quantify the angle *φ_a_*(*t*), a variety of measures from selective attention research combined with affective psychophysiology can be used, but it is critical to be able to link events of interest in a particular reality and modality (whether these events are strictly sensory, or rather significant from a more narrative or otherwise internal perspective) to measurable physiological events. This may seem trivial when considering more traditional experimental contexts in which the two “realities” across which attention may be divided are relatively simple (i.e. a mostly static visual scene containing two competing objects). However, this task becomes markedly more difficult when considering more advanced scenarios such as when the induced reality is emulating properties of an actual (physical) reality (e.g., in a 3D or 4D movie clip with immersive sound), or cases of augmented reality in which the induced stimuli are presented in direct competition with the full richness of the real world in which the participant is completely engaged. In such cases, the identification or annotation ‘by hand’ of physiologically relevant events in each reality for assessing degree of immersion can be impractical or even impossible, but recent computational efforts in machine learning, computer vision, and affective computing might help to identify/anntoate such events in the sensory streams automatically, see Eyben et al. [60], Hausfeld et al. [103], Trinh Van et al. [61], Höfling et al. [104], Flotho et al. [62].

#### Closed Loop Systems

While we have focused here on the use of physiological measures to assess the degree of immersion or perhaps the effectiveness of particular immersive methods, it is also possible to conceive of systems that use such measures as part of the immediate construction and ongoing maintenance of an induced reality. For example, virtual reality body suits (e.g., the Teslasuit, VR Electronics Ltd, London, UK) already incorporate physioloigcal measures from the wearer that could be used to modulate stimulus delivery in a contextually–specific manner. In applications in psychotherapy, adaptive VR can already be used to modulate the degree of stress imposed by a virtual scene (e.g. the height or narrowness of a virtual walkway above a pit in exposure therapy for fear of heights) [105]. Similarly, in applications in which modulating the degree of immersion might be important for the rehabilitation outcome, e.g., see Georgiev et al. [106], the proposed model could support close–loop systems to keep the immersion at the intended level, e.g., by sending an appropriate stimuli when a drift of attention away from the induced reality is detected (significant reduction of the angle *φ_a_*(*t*)), but perhaps also reducing properties conducive to immersion if the level of anxiety is identified as being too high.

### 4.3 Multimodal Integration

If we are to apply this model to understanding immersion in the context of existing, let alone future, virtual reality technologies, we must consider the case when multiple sense are stimulated to evoke the induced reality. In this article, we have remained vague about the specific nature of the stimuli in question, and have simply mapped abstract sensory information streams in the model which might or might not be induced by any one of several senses. However, multimodal integration likely plays a significant role in immersion, especially in technology driven induced reality settings.

A few decades ago, neuroscientists studied the world one sense at a time. Visual neuroscientists would study the world from a visual perspective and auditory neuroscientists would study the world from an auditory perspective. More recently, cognitive neuroscientists have realized that no sense operates in a vacuum. Instead, all senses interact in a process known as multisensory integration [107, 108, 58]. Indeed, Stein and Meredith [109] argued that there is no animal in which there is known to be a complete segregation of sensory processing. Several senses can summate to produce superadditive signals in the brain where 1+1=3. Alternatively, if the senses add up in the wrong way, they can produce subadditive integration and the senses can detract from each other. This depends on factors such as the relative timing, spatial coincidence, and semantic congruency (e.g., pairing a duck quack with a picture of a dog) of the different unimodal sensory stimuli in the multisensory perception. Stein and Meredith [109] said that “Integrated sensory inputs provide far richer experiences than would be predicted from their simple coexistence or the linear sum of their individual products.” In a similar line to our discussion in Sec. 4.1, imagine Alfred Hitchcock ‘Psycho’ without Bernard Herrmann’s music or several of the latest hit movies without Hans Zimmer’s compositions.

Based on the above principles and other principles such as sensory dominance, it may eventually be possible to determine the ideal integration of multisensory information for the strongest perceptual experience by considering stimulus aspects such as sensory modality, spatial location, relative timing, and other congruencies or incongruencies. In general, especially for the purposes of attentional capture, vision dominates the other senses, and it has been said that 70 % of attentional capture is based on vision, 20 % on audition and 10 % the rest, see Zimmerman [110]. However, even when multiple sensory signals are equally accessible, which sense dominates in a particular context or task depends on many factors, including perhaps their different ecological roles. For example, as discussed, e.g., by Francis [111], the human auditory system seems to have evolved to be particularly suited to the function of alerting, being sensitive to the occurrence of events in the environment over large distances and essentially in all directions [112], serving to quickly evaluate their relevance for subsequent action [113], and guiding action (including the direction of vision) appropriately, see Arnott and Alain [114]. In addition, audition can be more reliable for judging relative timing, and under certain circumstances can even overrule visual information, for example, creating two visual blinks out of one in the cross-modal double flash illusion (see Shams et al. [115]). Thus, competition between stimuli across different realities and different modalities can be quite complex, and further research will be necessary to determine how, or to what degree, information presented in different modalities can facilitate or interfere with the development and maintenance of immersion.

A variety of factors influence multisensory integration. For example, Spence and Santangelo [107] showed that the distance of the multisensory stimulus from the observer is important with the brain expecting an audiovisual stimulus that is far away (e.g., lightning and thunder) to have the visual aspect arrive first, while this expected gap becomes smaller when the observer is closer to the stimuli. Similarly, Marucci et al. [59] showed that task load matters for multisensory perception. When task load is high having combined visual, auditory, and tactile stimuli led to improved performance and an increased sense of immersion in the virtual environment compared to visual stimulation alone. For these reasons much effort has been devoted to integrating more senses into the VR experience. VR systems have been built that integrate vision, touch, audition, and smell (e.g., Sensync, Panagiotakopoulos et al. [3]) to produce optimally immersive environments. However, care must be taken to appropriately coordinate the information presented in each modality, in order to minimize conflicting cues and improve fidelity across modalities.

Thus, fidelity between multisensory stimuli is important and seems to enhance immersion. Thus, factors such as timing, spatial coincidence and semantic congruency play a large role for the immersive experience. Perceptually, audiovisual speech is typically far more understandable than audio or visual alone, but even relatively small asynchrony between visual (e.g. lip movements) and audio (speech sounds) signals can not only eliminate, but actually reverse this benefit [116, 117]. Semantically, a quacking dog might makes us instinctively react and snaps us out of the immersion. Thus, it seems like our brain has a particularly strong reaction to incongruencies to our expectations. This mechanism could potentially carry over to other types of rules as well such as when our expectations for multisensory stimuli are violated. Integrating these concepts in the two–competitor model could be an interesting line of future research.

### 4.4 Limitations

The presented model is necessarily very preliminary, and much work remains to be done for both, to further elucidate the model and to assess the validity of certain predictions made based on it.

#### Specific Immersion Settings

The current two–competitor model is very general as our intention was to provide a model of the role of attention in immersion that was broad enough to cover, at least preliminarily, circumstances as simple as reading a book to far more technologically advanced settings using augmented and virtual reality technology. Because of this, certain scenarios have not been analyzed in depth. For instance, our discussion concentrates on the state of immersion in the induced reality *per se*, and we disregard the “wow-effect” likely to be associated with using a new advanced technology and digital presence for the first time. Moreover, we have ignored unique properties of specific technologies, e.g., certain virtual reality headsets may be designed to physically block exogenous stimulation from the actual reality. As currently written, the model is intended to highlight the importance of competition between the two realities (induced and actual) for exogenous and endogenous attention, and we have also emphasized the essential role of internal attention to competing streams or objects as a potential source of distraction. Future research could absolutely investigate the development of the transfer functions Φ**_p_**(**s**(*t*)) and Ψ**_p_**(**s**(*t*)) for specific setting, e.g., reading, watching movies, or playing computer games, in fully virtual, augmented, or mixed realities as desired. We hope that the suggested two–competitor model paves the way in such directions by helping to specify the problems that arise by allowing them to be reduced to general functionalities mapped by explicit mathematical functions.

#### Interactivity and Motor Action

Technological approaches to induced realities allow for interactivity and motor action, a topic that has been studied in virtual reality based learning [118] using the Cognitive Affective Model of Immersive Learning (CAMIL), see Makransky and Petersen [6]. Moreover, the interplay of attention and motor action as such is an active field of current research, see Song [119] and some recent theories of attention have begun to suggest that a major role of attention is to subserve planning for action [120, 121]. Indeed, if we consider that “the purpose of the human brain is to use sensory representations to determine future actions” [122] it seems likely, then, that one way to increase the engagement of attention and hence immersion with an induced reality would be to increase (physical) interactivity with it as well [123].

#### Limits of a Two–Competitor Approach

Finally, in this article we have attempted to reconcile terminology and phrasing used in research on virtual, augmented, and mixed reality, which we have subsumed as induced reality, with that used in traditional attention research. This reconciliation has limits. For instance, we have tended to use a very broad conceptualization of immersion, to the extent that it would fit out definition to say that, when reading your tax report instead of a book, you might be ‘immersed’ in the sense that we have used here, i.e., paying attention to such a high degree that other stimuli are excluded. We might even expect that this state would exhibit many of the affective physiological states associated with such immersion, and yet the phrase ‘induced reality’ or ‘virtual reality’ (which we have tended to treat as the thing one is *immersed in*) might, for most of us unfortunately, not fit here. That is, it would seem incongruent to talk about being ‘immersed in the induced reality’ of one’s tax report. Thus the suggested two–competitor model just maps the case in which an induced reality is competing with an actual reality. Ultimately, we believe that this problem returns us to the deeper questions of how or to what degree ‘immersion’ must be distinguished from simple selective attention, on the one hand, and more complex and cognitively engaged phenomena such as ‘presence,’ ‘transportation,’ or ‘flow,’ on the other (see Agrawal et al. [16]). We have suggested that the role of internal attention is key to investigating this question, but further research is clearly necessary to provide a clearer picture.

## 5 Conclusions

We have analyzed the role of attention in immersion based on the hypothesis that attention is a necessary but not sufficient condition for immersion. By utilizing different modes of attention, we have developed the two–competitor model.

In this model, an induced and the actual reality are represented as orthogonal dimensions competing for the projection of attention. The two–competitor model allows for a quantitative implementation with an easy graphical interpretation, and helps to highlight the important link between different modes of attention in studying immersion. Even though the two–competitor model is a preliminary conceptual approach to study immersion, it is intended to provoke new, more specific research questions related to the role of attention in immersion and its interaction with affect and neurophysiological signals. In addition, we hope that it also stimulates researchers in cognitive and affective neuroscience to apply their knowledge to immersion research.

## Footnotes

1 The entire model implemented in Matlab2022b/SIMULINK, MathWorks, Inc, Natick, MA, USA. It will be publicly available at github after acceptance. Note that the absolute values in the distress transfer function were adjusted for the current setting as compared to Schneider et al. [57]).

## Acknowledgments

The first author is partially supported by the German Federal Ministry of Education and Research, Grant 13FH050KX1. The authors would like to thank Kyra GauthierDickey and David Thinnes for early discussions and technical support. The first author would like to thank also Steven A. Hillyard for his helpful feedback to some early thoughts on the topic.

